# Embracing fine-root system complexity to improve the predictive understanding of ecosystem functioning

**DOI:** 10.1101/2022.10.07.511037

**Authors:** Bin Wang, M. Luke McCormack, Daniel M. Ricciuto, Xiaojuan Yang, Colleen M. Iversen

## Abstract

Projecting the functioning of the biosphere requires a holistic consideration of whole-ecosystem processes. Although improving leaf and canopy processes has been the focus of ecosystem model development since the 1970s, the arbitrary homogenization of fine-root systems into a single pool is at odds with observations. This discrepancy has increased in the last two decades as accelerated conceptual and empirical advances have revealed functional differentiation and cooperation conferred by the hierarchical structure of fine-root orders and associations with mycorrhizal fungi in fine-root systems. To close this model-data gap, we propose a 3-pool structure comprising Transport and Absorptive fine roots with Mycorrhizal fungi (TAM) to model vertically resolved fine-root systems across organizational and spatial-temporal scales. A comparison of TAM to the single fine-root structure in a state-of-the-art Earth System Model using the ‘big-leaf’ approach demonstrates robust impacts on carbon cycling in temperate forests, lending further quantitative support to the empirical and theoretical basis for TAM. Strong support in both theory and practice therefore suggests a move beyond the useful but incorrect paradigm of single-pool homogenization, echoing a broad trend of embracing ecological complexities in terrestrial ecosystem modelling. Although challenges lay ahead towards realizing TAM in ecologically realistic demography models simulating emergent functioning from pattern and diversity, adoption of TAM by both modelers and empiricists holds promise to build a better predictive understanding of ecosystem functioning in the context of global change.

## 1. Introduction

Earth System Models (ESMs) grapple with low confidence in projecting the functioning of the biosphere within the Earth System across space and over time. Many efforts have been made to improve predictability regarding the structure and functions of terrestrial ecosystems since the 1970s, though this work has mostly focused on the aboveground leaf and canopy processes (e.g., **Sinclair et al., 1976; Friend et al. 2014**; **Lovenduski and Bonan 2017**). Meanwhile, plant roots have been integral to the evolution of plant form and function (e.g., **Raven and Edwards 2001**) and sit at the nexus of plant-microbe-soil interactions in the biosphere, linking the atmosphere and pedosphere (e.g., **Bardgett et al. 2014; Freschet et al. 2021**). However, current models treat fine-root systems—the ephemeral portion of the root system responsible for water and nutrient acquisition—rudimentarily, and more importantly, lag behind empirical and theoretical advances with regard to fine-root system complexity (e.g., **Fitter 1982; Pregitzer 2002; Phillips et al. 2013; McCormack et al. 2015a**). This model-data discrepancy likely contributes to the poor representation of terrestrial ecosystem feedbacks to increasing CO2 within ESMs (e.g., **Warren et al. 2014; Bonan and Doney 2018**). Therefore, embracing fine-root system complexity to improve structural realism is expected to increase the prognostic capability of terrestrial ecosystem models in biosphere-atmosphere interactions.

Over the past two decades belowground ecologists have made tremendous progress in revealing the heterogeneity within fine-root systems. Within a fine-root system, plants have evolved roots with unique morphological, anatomical, chemical, and physiological features that differ among root branching orders (e.g., **Pregitzer et al. 2002; Atucha et al. 2021**), as well as varying microbial associations with both mycorrhizal fungi and bacteria (e.g., **Sen and Jenik 1962; King et al. 2021**). These structural differences support both functional differentiation and cooperation within fine-root systems; fine roots of higher orders are primarily responsible for transport of water and nutrients to the rest of the plant, while finer and more distal roots in cooperation with mycorrhizal fungi are responsible for nutrient and water absorption (e.g., **Hishi & Takeda 2005; McCormack et al. 2015a; Wang et al. 2022**). This heterogeneity, and degree of coordination with aboveground plant activities, changes with species and habitat, forming diverse whole-plant strategies across biomes under physical and functional constraints (e.g., **McCormack and Iversen 2019; Weigelt et al. 2021)**.

By contrast, fine-root systems in ecosystem models are often treated without accounting for their structural and functional heterogeneity. Currently, the prevailing option in terrestrial ecosystem modelling remains a single fine-root pool (e.g., **Kleidon & Heimann 1998**; **Zeng 2001; Warren et al. 2015; Burrows et al. 2020**). Although empiricists have advocated for specifically representing arbuscular mycorrhizal fungi traits and function (**Treseder 2016**) and modelers have even already incorporated impacts of mycorrhizal fungi into models either implicitly (e.g., **Woodward and Smith 1994; Brzostek et al. 2014**) or explicitly (e.g., **Hunt et al. 1991; Orwin et al. 2011; Sulman et al. 2018**), no efforts yet holistically integrate the complexity of fine-root system with a balanced perspective of both fine roots and mycorrhizal fungi. This oversimplification of fine-root system structure fundamentally limits accurate representation of its functioning and hence above- and below-ground interactions (e.g., **Smithwick et al. 2014; Warren et al. 2015**). In addition, such a simplification forms an asynchronous development in terrestrial ecosystem models relative to other components (notably photosynthesis and canopy processes) that likely imparts a structural limit on improvements to whole model performance. For instance, fine-root dynamics are normally assumed to follow leaf phenology closely (e.g., **Burrows et al. 2020**). Of course, there are individual-level plant models, especially for crops, simulating explicit root structures (e.g., **Pointurier et al. 2021**). However, it has been a challenge to integrate theoretical and empirical advances at scales relevant for ecosystem models while balancing model simplicity and realism.

Such a large model-theory discrepancy raises the question of whether models can improve performance with an appreciation of fine-root system complexity. The core of this question about a single pool versus a more explicit treatment, by analogy, mirrors the debates in vegetation modelling on one layer versus multilayer canopy modelling (e.g., **Raupach and Finnigan 1988**), on individual-based versus various functional group modelling (e.g., **Smith et al. 1997; Shugart et al. 2018**), and on different lumping approaches in modelling various complex physical, chemical, and biological systems more generally (e.g., **Okino and Mavrovouniotis 1998)**. For example, comparisons of one- versus multi-layer canopy simulations have led to the conclusion that land surface models should move beyond the useful but incorrect paradigm of single-layer canopy scheme to help resolve uncertainties in canopy processes (**Bonan et al. 2021**). Similarly, current models show large uncertainties related to belowground processes; for example, a comprehensive sensitivity analysis of the ELM (**Box 1**) revealed large prediction uncertainty arising from uncertainty in root-related parameters and processes (**Ricciuto et al. 2018**). Constraining model uncertainty requires a model structure that can predict measurable quantities in real fine-root systems. Furthermore, improving the representation of fine roots may help model performance by identifying other coupled but deficient components. Therefore, dedicated efforts are warranted to explore alternatives to the single fine-root pool in current terrestrial ecosystem models. Meanwhile, the increasingly available data on fine-root traits (FRED; **Iversen et al. 2017; Box 1**) and fungal traits (e.g., **Zanne et al. 2020; Iversen and McCormack 2021**) can facilitate these efforts.

To this end, we propose a generalized 3-pool structure including Transport and Absorptive fine roots as well as Mycorrhizal fungi (TAM) to represent fine-root systems across organizational and spatio-temporal scales in terrestrial ecosystem models. This function-based TAM structure is intended to approximate the high-dimensional heterogeneity within the hierarchical branching fine-root systems and to serve as an explicit but tractable approach to model fine-root system functioning. The overarching objective of this viewpoint is to argue for adoption of this structure as the quantitative keystone of the bridge between modelers and empiricists. To achieve this objective, we evaluate this TAM structure both theoretically and quantitatively. After elaborating on the conceptual, theoretical, and empirical bases of TAM, we specifically address the following three questions: First, how to realize TAM in ecosystem models? Second, how does this new structure alter simulations of major ecosystem carbon fluxes (e.g., productivity and soil respiration) and pools (e.g., fine-root biomass and soil carbon storage)? Third, what are the uncertainties and challenges in broadly adopting this TAM structure?

We answer the first question by proposing a high-level framework of carbon partitioning, the phenological patterns of root and fungal birth, growth, and death, and the vertical distribution of fine roots and fungi throughout the soil profile that can be integrated into existing modelling paradigms of varying complexity in terms of vegetation structure. Starting with the relatively simple big-leaf modeling paradigm, we then answer the second question with an example implementation in the context of a representative big-leaf model with a single fine-root pool— ELM—in two different temperate forest types (deciduous and evergreen forests) across the Eastern United States. For a proof-of-concept purpose the focus of this example is mostly descriptive rather than analytical. By showing significant impacts of TAM in a relatively simple model structure we do not fully address the question of whether TAM really improves model predictability. Rather, we believe a combination of the conceptual, theoretical, and empirical bases and an initial demonstration in this viewpoint lays the groundwork for broad adoption and evaluation of TAM in different modelling paradigms against observations at varying scales. Accordingly, we answer the third question by discussing uncertainties and challenges to fully exploit its potential. We conclude by pointing out potential implications of TAM for guiding empirical research, understanding ecosystem functioning, and improving ESMs prognosis.

#### Box 1 Terminology

**Fine-root system**: Root-mycorrhizal association at the individual plant level comprising fine roots of different orders and associated mycorrhizal fungi.

**TAM**: An approach argued for in this viewpoint to abstract fine-root systems with two pools representing the traits and function of fine roots—transport roots (T) and absorptive roots (A)—as well as mycorrhizal fungi (M).

**FRED (Fine Root Ecology Database)**: a database conceived and built by **Iversen et al. (2017)** that houses fine-root trait observations from across the world (https://roots.ornl.gov/).

**ESM (Earth System Model)**: A coupled-climate model that explicitly simulates the atmosphere, the ocean, and the land surface in the Earth system, of which the land component is usually referred to as the land model.

**Terrestrial Ecosystem Model**: Models that simulate ecosystems across spatial-temporal scales in the terrestrial biosphere, which is synonymous with land model, terrestrial biosphere model, or ecosystem model in this study.

**Big-leaf model**: Terrestrial ecosystem models that represent vegetation using prescribed, static fractional coverage of different Plant Functional Types (an approach to aggregate and simplify global plant diversity) with either a single canopy layer (i.e., one big-leaf model) or sun and shaded layers (i.e., two big-leaf model).

**ELM**: The land model using the big-leaf approach with a single fine-root pool in the Energy Exascale Earth System Model (E3SM), a state-of-the-art ESM developed by the US Department of Energy (**Golaz et al. 2019)**.

**Demography model**: Terrestrial ecosystem models that can simulate dynamics of vegetation pattern (e.g., forested mosaic of gaps) with demographic processes of growth, mortality, and reproduction.

**Demand-driven approach**: An approach for formulating nutrient uptake by fine-root systems of plants or heterotrophic microbes based exclusively on their nutrient demands determined by their potential growth or decomposition rates. The competition between plants and microbes for nutrients is resolved by comparing their demands, which is thus referred to as relative-demand approach.

**Trait divergence**: Trait variability or heterogeneity where a 3-pool TAM model allows for unique trait values for each pool while a 1-pool fine-root model only allows a single value to represent the fine-root system.

**Preservation**: A procedure to keep the same whole bulk fine-root system property (e.g., C/N in this study) while capturing trait divergence from a 1-pool to 3-pool TAM structure.

**Partitioning**: The division of plant photosynthates allocated to one of the TAM pools. **Distribution**: The explicit vertical representation of photosynthates in a soil profile partitioned to one of the TAM pools.

## 2. Conceptual, theoretical, and empirical basis of the TAM structure

TAM arises from a conceptual shift of fine-root systems. Fine-root systems develop structural and functional differentiation within an individual plant and express substantial variation among individuals, across species, and across biomes (e.g., **Weigelt et al. 2021**). Structurally, fine-root systems bear a hierarchical branching structure with different orders; each order displays different properties in its anatomy, morphology, chemistry, physiology, lifespan, and fungal colonization (e.g., **Pregitzer et al. 2002**; **Guo et al., 2008; McCormack et al., 2017; Klimešová & Herben 2021; Zhou et al. 2022**). This structural variation underlies distinct functional differentiation within fine-root systems. Therefore, an arbitrary homogenization of such differentiation below the 2-mm diameter threshold as functionally equivalent fine roots is problematic. Instead, a new conceptualization of functional differentiation is argued in terms of transport (coarser, high-order fine roots), absorption (finer, low-order fine roots), and facilitation by colonizing mycorrhizal fungi (**McCormack et al 2015a**). Therefore, following a long tradition of function-based lumping of objects in both empirical and quantitative ecology (e.g., **Raunkiaer 1934; Root 1967; Grime 1974; Smith et al. 1997**), we aggregate fine-root systems of different orders and associations with mycorrhizal fungi into three functional pools that comprise transport roots, absorptive roots, and mycorrhizal fungi—TAM (**Fig.1**).

**Fig. 1.**
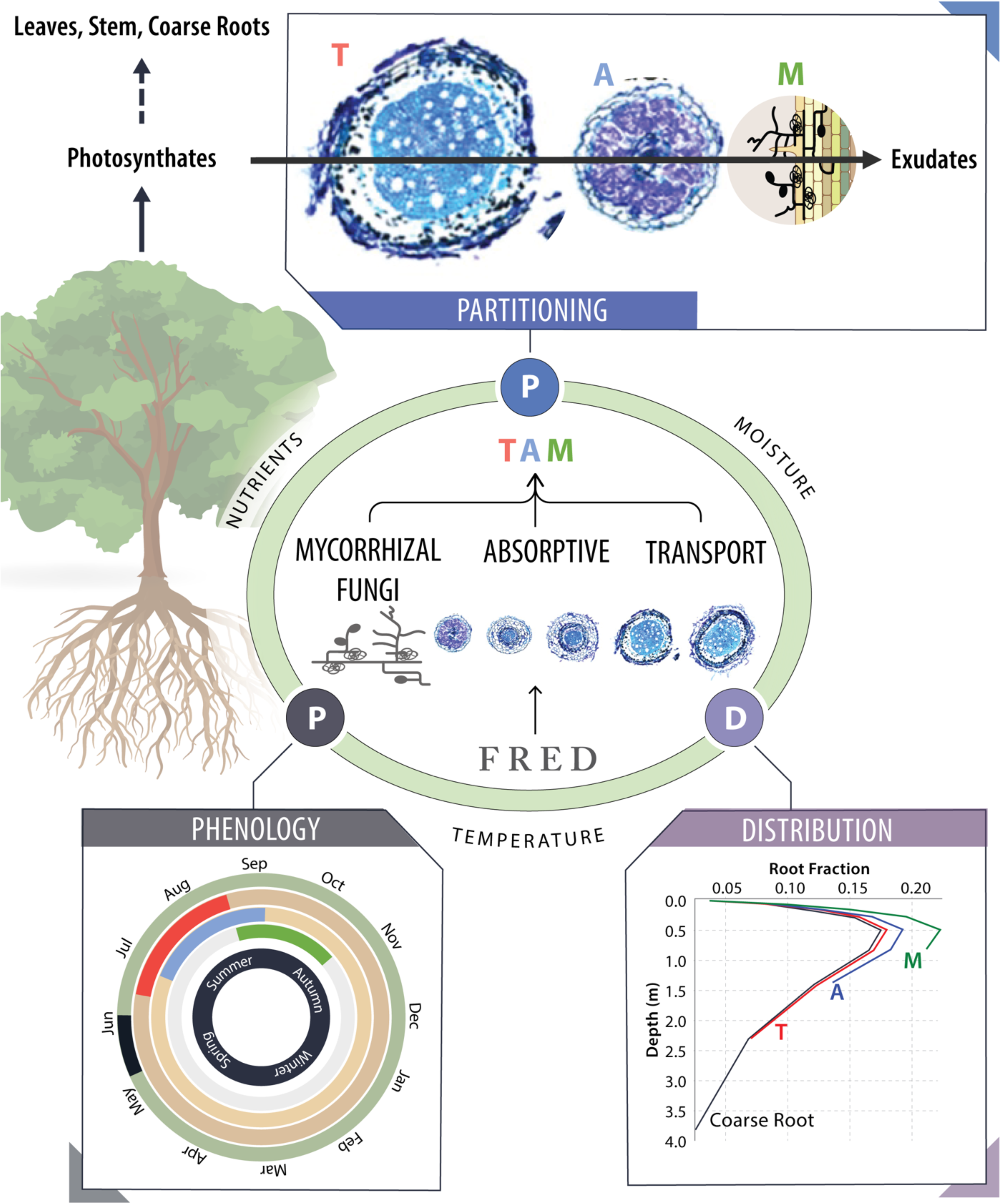
Schematic of the 3-pool TAM structure governed by interrelated processes of partitioning, phenology, and distribution under environmental constraints. TAM is conceived to be informed by order- or function-based explicit measurements of fine-root and mycorrhizal fungal traits as compiled in various databases, e.g., FRED. Partitioning of photosynthates to the outside of fine-root systems as exudates is also indicated, though not yet explicitly treated in the current study. Asynchronous phenology within TAM and with leaf phenology is illustrated with approximate periods of production (indicated by colored bands) for leaves, T, A, and M pools based on a *Quercus alba* Plot at The Morton Arboretum, USA in 2019. In addition, the different vertical distribution of TAM (in reference to the coarse root distribution) is for an illustrative purpose only; distribution of new carbon still assumed to be the same across TAM pools though with different turnover rates. The cross sections of 5 fine-root orders are reproduced from McCormack et al. (2015a).

Theoretically, this 3-pool TAM structure allows a balanced perspective of fine roots and fungi in ecosystem models while capturing both functional differentiation and cooperation within the fine-root systems. A single fine-root pool oversimplifies fine-root systems and misses important symbiotic associations with mycorrhizal fungi. Recognizing the importance of mycorrhizal fungi, some models have accounted for these fine-root partners as an implicit component (e.g., **Koide and Elliot 1989; Kirschbaum and Paul 2002**; **Brzostek et al. 2014**). While these efforts have contributed to more accurate simulations of vegetation nutrient uptake and soil organic matter decomposition in general, these plant-centric models do not include an active mycorrhizal pool that enables a balanced perspective of plant and mycorrhizal fungi. However, even when an explicit mycorrhizal fungal pool is included (e.g., **Hunt et al. 1991; Orwin et al. 2011; Sulman et al. 2019**), a homogenous treatment of fine roots still misses the structural and functional differentiation arising from fine-root orders (**Fig.1**). Of course, a discretized 3-pool structure cannot fully capture the high-dimensional fine-root system heterogeneity (typically five or more root orders plus mycorrhizal fungal colonization, e.g., **Pregitzer et al. 2002; Guo et al. 2008**). Individual-level plant models, especially for crops, may be suitable for an order-based continuous treatment (e.g., **Couvreur et al. 2012; Pointurier et al. 2021**), whereas ecosystem models often need to balance biological and ecological complexity with computational efficiency. Further increasing complexity by having more pools to account for each fine-root order would likely fall into the trap of diminishing returns in performance with an increasing parameterization challenge (e.g., **Transtrum et al. 2015**).

Parameterization of a 3-pool TAM structure has empirical support across organizational and spatial-temporal scales in the biosphere. TAM confers generality across scales by enabling adaptable fine-root pools. While T and A capture widespread functional differentiation between transport and absorption, they can be effectively reduced to one pool to accommodate rare species without such a clear differentiation (**McCormack et al. 2015a**). Also, the M pool, in practice, can be attributed to hyphal mycelium of arbuscular (AM), ectomycorrhizal (ECM), and/or ericoid fungi, depending on the specific site and/or geographic context (e.g., **Read 1991; Soudzilovskaia et al. 2015**). Besides this adaptability, the empirical support is facilitated by availability of trait databases. In the context of a belowground data revolution [see a synthesis by **Iversen and McCormack (2021**)], databases with increasing spatial coverage of explicit fine-root traits and fungal traits aggregated from either function- or order-based measurement make it feasible to parameterize TAM across scales in terrestrial ecosystem models. Notably, the Fine-Root Ecology Database (FRED), houses root trait observations from across c. 4600 unique plant species and continues to grow (**Iversen et al. 2017; Iversen and McCormack 2021**). Such an empirical support warrants realizing and testing TAM in terrestrial ecosystem models.

## 3. Realizing the TAM structure in terrestrial ecosystem models

To realize TAM in terrestrial ecosystem models, we propose a high-level framework of partitioning, phenology, and distribution to encapsulate processes towards simulating a temporally and vertically resolved 3-pool fine-root system (**Fig.1**). Partitioning is responsible for determining the magnitude of carbon allocated from recent photosynthates or storage to the T, A, and M pools, which is dynamic in nature arising from changes in biotic and abiotic environment from edaphic factors (e.g., nutrient availability) to atmospheric changes (e.g., elevated CO2) (**Mooney 1972; Chapin et al. 2009; Drigo et al. 2010; Gorka et al. 2019; Ouimette et al. 2020; Prescott et al. 2020**). Phenology then controls the timing of partitioning. Here, we are highlighting phenology as an essential component because of recurring observations of above- and below-ground phenological asynchronicity (e.g., **Steinaker et al. 2010; Abramoff & Finzi 2015**; **McCormack et al. 2015b; Radville et al. 2016; Iversen et al. 2018**). In addition to variations in phenology among the TAM structures, phenology varies with soil depth (e.g., **Maeght et al. 2015**). This vertical variability makes it essential to represent the vertical distribution within the soil profile of carbon and nutrients partitioned to TAM pools at each time-step (e.g., **Tumber-Dávila et al. 2022**). Moreover, traits of fine roots and mycorrhizal fungi vary vertically (e.g., **McElrone et al. 2004; Robin et al. 2019**). This vertical variability feeds back to influence partitioning. Therefore, these three components are interrelated in such a way that requires a close coupling of dynamic partitioning with phenology temporally and distribution vertically. These components and their interactions can be integrated into existing models by coupling with upstream photosynthesis and allocation and with downstream soil biogeochemistry.

Starting this integration from a simple model structure is practical in the sense of observing impacts of TAM while paving the way for realization and evaluation in more complicated model structures (**Fig.2**). Existing models, built with different assumptions for different purposes, vary in their structural realism of representing vegetation pattern and process under three different paradigms. Integrating TAM into models under these paradigms can incorporate different organizational levels across spatial-temporal scales, which, though exciting, trades off with feasibility in terms of formulation, parameterization, and evaluation. Testing proof-of-concept impacts of TAM under the relatively simple big-leaf paradigm is expected to help identify both opportunities and challenges to realize TAM under structurally more realistic vegetation paradigms.

**Fig. 2.**
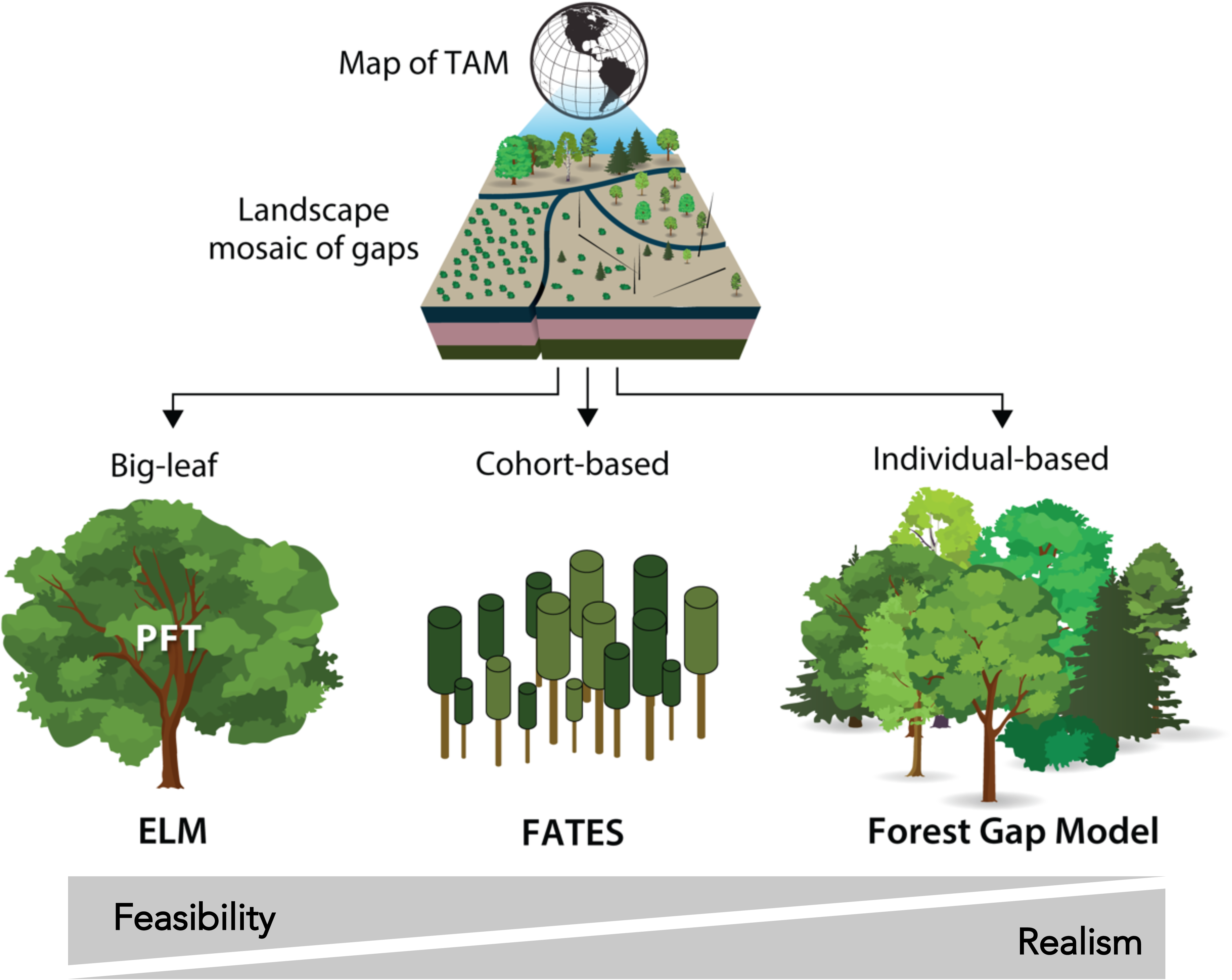
Realizing TAM with a feasibility-realism tradeoff in terrestrial ecosystem models under three paradigms of vegetation structure. The landscape of vegetation, particularly forests, is a mosaic of gaps (or patches) at different successional stages under exogeneous and endogenous disturbances, structuring vegetation dynamics both vertically by size and horizontally by age (e.g., Watt 1947; Bormann and Likens 1979). The three paradigms capture such pattern and process of vegetation to different extents. Big-leaf models (e.g., ELM), representing global vegetation in fractional coverage using a few PFTs, do not capture local plant diversity and cannot predict structural dynamics. By contrast, in capturing the vertical and spatial structure of forests, individual-based forest gap models explicitly simulate growth, mortality, and reproduction of very single individual in an array of gaps (e.g., Shugart 1984; Shugart et al. 2018). By approximating the spatial and vertical structuring of vegetation without an explicit parameterization of different individuals (e.g., Kohyama 1993), cohort-based models [e.g., FATES: Functionally Assembled Terrestrial Ecosystem Simulator (Fisher et al. 2018) and ED: Ecosystem Demography model (Moorcroft et al. 2001)] simplify individual-based gap models by utilizing cohorts of like-sized plants under the traditional PFT scheme. Models under these paradigms assume a single fine-root pool; realizing TAM in these models can capture different organizational levels across spatio- temporal scales in the biosphere. ELM, as a representative of the relatively simplest big-leaf model, is taken as an example for TAM demonstration in this viewpoint.

Therefore, we examine impacts of TAM in the context of the state-of-the-art Energy Exascale Earth System Model Land Model, ELM. ELM is a typical big-leaf model with a single fine-root pool based on the traditional scheme of representing global plant diversity using relatively few PFTs, which has been extensively used for land-atmosphere interactions regionally and globally (e.g., **Golaz et al. 2019; Burrows et al. 2020**). Moreover, ELM uses the relative- demand approach as a default configuration for plant-microbe nutrient competition (**Box 1**), which does not require an explicit representation of fine-root and fungal functions and remains a prevailing approach in earth system modelling as a simplification of plant-microbe competition for nutrients (e.g., **Thornton et al. 2007; Yang et al. 2014**). Such formulations of ELM make it feasible for building and testing the TAM structure for a proof-of-concept purpose.

## 4. Impacts of TAM on forest ecosystems: an example realization

We introduced a series of specific changes within the above framework to realize a 3-pool TAM structure and examined its impacts against the 1-pool structure in ELM. In contrast to the constraint of a single-pool structure, TAM allows trait divergence (**Box 1**) among the fine-root and fungal pools with respect to carbon to nitrogen ratio (C/N), longevity, and chemical composition, as well as respiration (**Table 1**), following observed trait variability within fine-root systems [see the synthesis of **McCormack et al. (2015a**)]. To partition the carbon fixed by a plant to each of the TAM pools, we introduced three parameters—fractions of the total allocation to a fine-root system determined by the prevailing assumption of a 1:1 allometry between leaf and fine-root system. These fractions together with C/N of the three TAM pools determine the C/N of bulk fine-root system (**Eq. 1** in the **Appendix**), which, in combination with other structural tissues (leaf, stem, and coarse-root), determine whole-plant stoichiometry and thus dictate nitrogen demand and uptake under the demand-driven approach. We then decoupled the timing of partitioning to TAM pools from leaf phenology, enabling an independent control on initiation (as a function of growing degree days) and turnover (determined by prescribed longevity) of TAM for both deciduous and evergreen PFTs. The turnover is further constrained by a depth-correction term to capture the widely observed pattern of decreasing turnover with soil depth (e.g., **Baddeley and Waston 2005; McCormack et al. 2012; Gu et al. 2017**). To vertically distribute the carbon partitioned to TAM pools within the soil profile we introduced a formulation of dynamic distribution based on changing profile of nutrient availability instead of the prevailing static assumption (e.g., **Zeng 2001; Drewniak 2019**). This nutrient-based dynamic distribution allows more of the partitioning to go to the depth where the relative availability of nitrogen is higher. We implemented these changes to different extents to examine structural uncertainty while accounting for parameter uncertainty in a deciduous forest and an evergreen forest in the temperate forest biome of North America only for an illustration of impacts of TAM against ELM (see **Methods** in the **Appendix** for details).

**Table 1.** Measured traits used to parameterize TAM in the example realization and expected to be used to directly parameterize a functionally explicit TAM in terrestrial ecosystem models.

| Category | Trait | T | A | M | Database |
| --- | --- | --- | --- | --- | --- |
| chemistry | **stoichiometry** |  |  |  | FRED |
|  | **chemical composition** |  |  |  | FRED |
| demography | **longevity** |  |  |  | FRED |
|  | **phenology** |  |  |  |  |
| morphology | diameter |  |  |  | FRED |
|  | tissue density |  |  |  | FRED |
|  | specific root length |  |  |  | FRED |
| physiology/metabolism | **respiration rate** |  |  |  | FRED |
|  | **temperature sensitivity of respiration ($Q_{10}$)** |  |  |  |  |
|  | transporter enzyme production rate |  |  |  |  |
|  | exoenzyme production rate |  |  |  |  |
| nutrient acquisition | transporter enzyme kinetics ($K_m$ & $V_{max}$) | | | | |
| | exoenzyme degradation kinetics ($K_m$ & $V_{max}$) | | | | |
| water uptake | hydraulic conductivity |  |  |  | FRED |
|  | water uptake |  |  |  |  |
| profile | max rooting depth |  |  |  | FRED, RSIP |

Note that the traits listed above are those that can directly inform TAM parameterization. Many of the measurements listed in databases (e.g., FRED) are emergent properties and could instead be used for model validation. Bolded traits are used for demonstration in this viewpoint, while grey cells indicate applicability of the traits to TAM. Database indicates coverage of the traits by FRED (which does not necessarily mean a full coverage of all TAM structures) or not (which clearly suggest targets of more empirical efforts). RSIP (Root Systems of Individual Plants) database is a primary source for root system size. See **Iversen and McCormack (2022)** for a synthesis of available databases of fine-root and mycorrhizal fungi traits.

### 4.1 TAM impacts under preservation of bulk fine-root stoichiometry

First, changing the structure from 1 to 3 pools but preserving the C/N of bulk fine-root system can give us a parsimonious view of impacts arising from trait divergence in TAM. A direct reason of preservation (**Box 1**) is that C/N of the bulk fine-root system, in principle, should be the same irrespective of the number of pools used in model representation. Keeping the same C/N is also useful in the sense of using existing field observations of bulk fine-root C/N to constrain the model. More importantly, with a demand-driven approach in ELM, C/N of the bulk fine-root system instead of C/N of any of the TAM components dictates plant nitrogen demand and uptake. Therefore, a preservation provides two benefits: it keeps the same nutrient uptake capability as the single-pool model while capturing the divergence in C/N among TAM pools. The preservation is achieved via pairing a set of C/N values with a set of partitioning fractions of TAM (see **Methods** in the **Appendix**). To capture the wide spectrum of plant partitioning variability among different belowground sinks, we consider two sets of partitioning fractions forming two contrasting partitioning patterns: a descending pattern (i.e., increasingly less carbon partitioned to the mycorrhizal fungi) and an ascending pattern (i.e., increasingly more carbon partitioned to the mycorrhizal fungi). Under both patterns, the tissue C/N decreases while turnover increases from T, to A, and then to the M pool; the M has the lowest C/N and fastest turnover, reflecting their consistently lower C/N and short lifespans (**Allen and Kitajima 2013; Zanne et al. 2020**). Under the same phenology, we compared TAM under the two partitioning patterns against ELM first by assuming the same fixed vertical distribution and then by adding dynamic vertical distribution contingent on relative nutrient availability (**Fig. 3**).

**Fig. 3.**
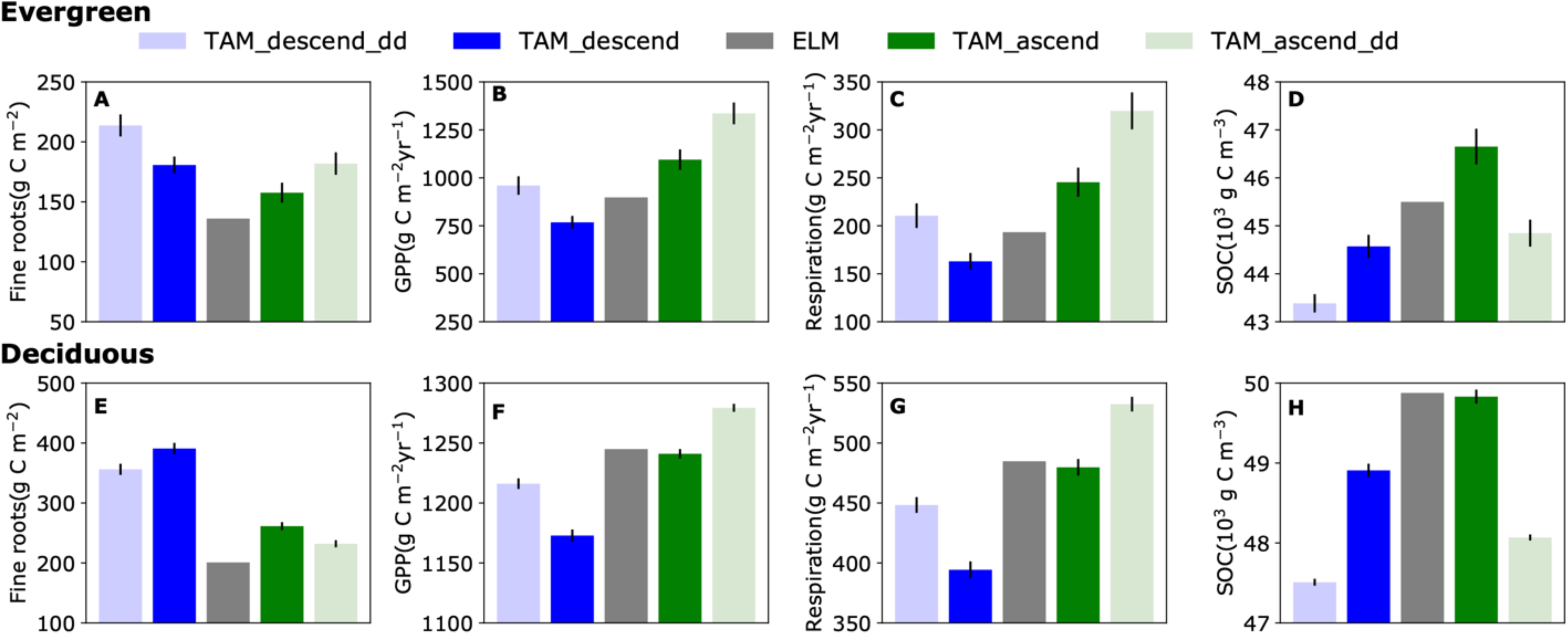
TAM impacts against the 1-pool fine-root model on temperate evergreen (A-D) and deciduous forests (E-H) under preservation of the bulk fine-root C/N. The preservation is differentiated between two partitioning patterns: TAM_descend (increasingly less partitioned to mycorrhizal fungi) and TAM_ascend (increasingly more partitioned to the mycorrhizal fungi), on top of which comparisons are made by further adding dynamic distribution (TAM_descend_dd and TAM_ascend_dd). Note these comparisons were made under an assumption of the same phenology between TAM and ELM. Fluxes of GPP (B, F) and heterotrophic respiration (C, G) are aggregated annual values, while pools of fine-root mass and SOC are mean daily values. The variance (95% confidence interval) arises from parameter uncertainty with respect to longevity and fine-root chemistry. Note different y-axis scales across panels are for distinguishing between different scenarios. See seasonal dynamics of GPP and fine roots, as well as leaf biomass in Supplementary Fig. 1 in the Appendix.

With the same static distribution, since both the stoichiometry and the phenology are the same, significant partitioning-dependent impacts of TAM compared against the 1-pool model can be attributed to changes in turnover and litter inputs arising from explicit low to high turnover rates and high to low C/N moving from T through M pools. The impacts are directly reflected in changes in fine-root biomass (including mycorrhizal hyphae biomass). When reaching equilibrium in both the evergreen and deciduous forest, under the descending partitioning pattern total fine-root biomass increased relative to the single-pool model owing to the lowest turnover rate of T roots (**Fig. 3A, E**). Similar, but somewhat dampened responses, were seen under the ascending pattern as there was relatively more carbon portioned to the M pool which has the highest turnover rate and resulted in less total standing biomass. These changes under equilibrium also came from feedbacks associated with changing temporal patterns of fine-root litter inputs and subsequent changes in GPP (**Fig. 3B, F**), soil heterotrophic respiration (**Fig. 3C, G**), and soil organic carbon storage (**Fig. 3D, H**), although these shifts were insignificant for the deciduous forest under the ascending pattern.

Enabling the dynamic vertical distribution further enhanced the effects of changing turnover of fine-root systems and hence litter inputs according to soil nutrient availability. Because this dynamic distribution increased inputs of fine-root and mycorrhizal fungal litters into soil layers where N availability was relatively higher, a reduction in N limitation led to increased soil respiration and GPP (**Fig. 3D, H**) but with a net effect of soil carbon stock decline under both patterns (**Fig. 3C, G**). However, fine-root biomass experienced a continued increase in the evergreen forest compared to a pause in the deciduous forest because relatively more biomasses accumulated in deeper soils with slower turnover rates in evergreen forest than in the deciduous forest (**Fig. 3D, H**). These new changes under dynamic distribution highlight the role of vertical variability of TAM in influencing plant-soil interactions. In short, even parsimonious TAM has important implications for carbon cycling under stoichiometry preservation while capturing observed differentiation within fine-root systems.

### 4.2 TAM dampens forest productivity and soil carbon storage

Implementing TAM without the constraint of stoichiometry preservation consistently resulted in reduced ecosystem productivity and soil carbon stock regardless of forest type compared to the 1-pool model (**Fig. 4**). With a full accounting of parameter uncertainty informed by observations (see **Methods** in **Appendix** for details), on average annual GPP was reduced by 20.3 % and 36.3 % while soil organic carbon was reduced by 5.9 % and 6.2 % in the evergreen and deciduous forests, respectively. These reductions in equilibrium originated from changing allocation (different C/N of fine-root systems) and explicit fine-root turnover, as indicated by the increased, though not seasonally consistent, fine-root biomass in equilibrium **(Fig. 4A, E)**. Over time these changes led to aggravated nutrient limitation (**Supplementary Fig. 2**), together with decreased leaf area, contributing to declines in GPP (**Fig. 4B, F**). Such a GPP reduction eventually resulted in decreased heterotrophic respiration and a net decline of soil organic carbon (**Fig. 4D, H**). These robust changes arising from TAM suggest that 1-pool models may overestimate forest productivity and soil carbon storage, which is in line with the general notion of a lack of accurate sink-limited growth in current terrestrial ecosystem models leading to an overestimation of land carbon sink (e.g., **Cabon et al. 2022**). In summary, the improved realism of TAM informed by empirical trait divergence can show robust impacts while accounting for parameter uncertainty.

**Fig. 4.**
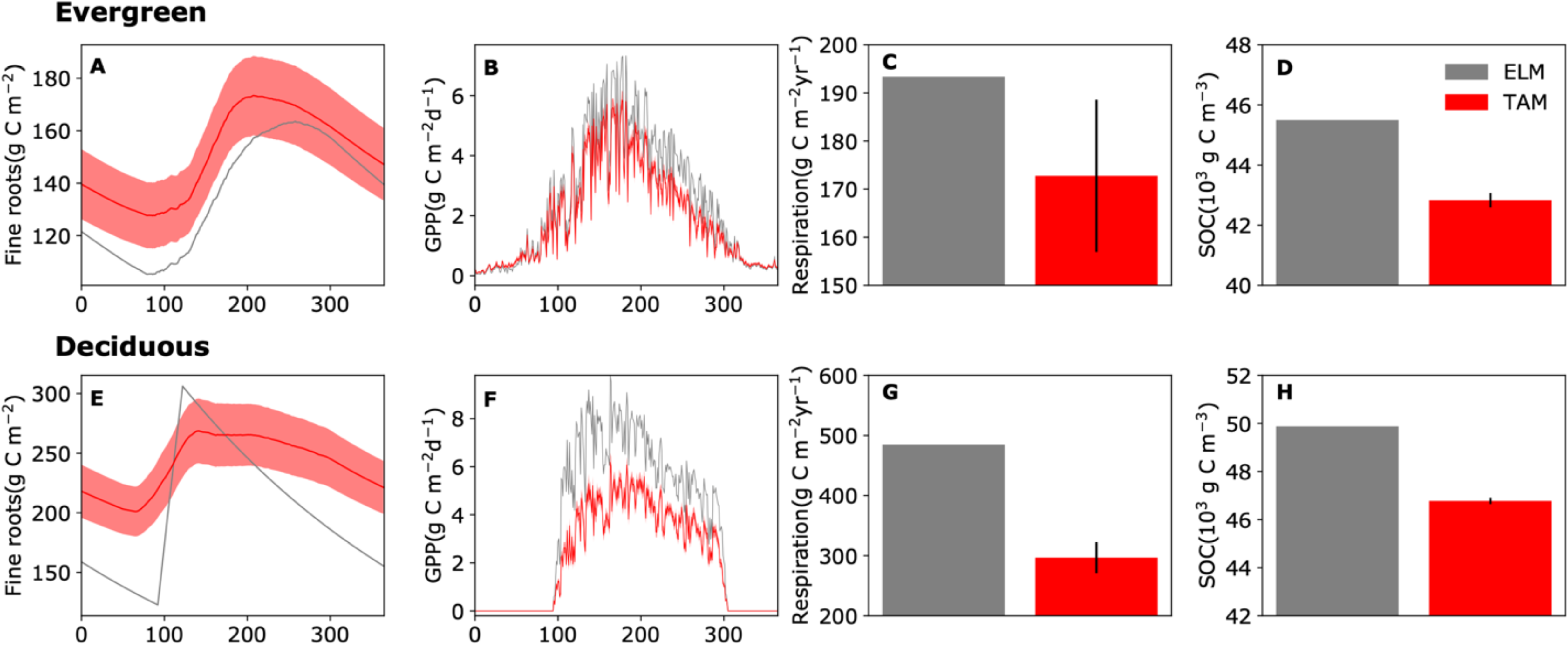
TAM impacts against the 1-pool fine-root model on temperate evergreen (A-D) and deciduous forests (E-H) without the restraint of preservation. Respiration is aggregated annual values (C, G) and SOC is averaged across a whole year (D, H). The variance (95% confidence interval) arose from accounting for all uncertain parameters introduced to realize TAM (see Supplementary Table 1).

### 4.3 TAM echoes embracing complexity in ecosystem modelling in general

These results provide strong quantitative support for TAM. The examples above demonstrate significant impacts of the explicit TAM structure against a homogeneous single-pool structure while accounting for structural and parameter uncertainty, especially considering the case of parsimonious stoichiometry preservation. Admittedly, this demonstration is based on only one simplified, though generally representative, vegetation modelling scheme, and it remains premature to claim that this TAM structure has improved model predictability without testing TAM across a broader range of sites in demography models (**Fig.2**). However, such a shift from implicit to explicit reflects a broad notion of embracing biological and ecological complexity in ecosystems from above- to below-ground to improve model performance. A notable aboveground shift in vegetation structure is from the big-leaf approach using a single-layer canopy to explicitly modeling multi-layer canopy structures (e.g., **Sinclair et al., 1976; Norman, 1993; Yan et al. 2017; Bonan et al. 2021**), and further to a paradigm simulating vegetation dynamics in the spatial and vertical structuring by age and size (e.g., **Shugart 1984; Kohyama 1993; Moorcroft et al. 2001; Wang et al. 2016; Shugart et al. 2018; Fisher et al. 2018**). A similar belowground example is the shift from simulating the tremendously complex soil microbiomes first implicitly (e.g., **Parton et al. 1987**), then as a ‘big microbe (e.g., **Allison et al. 2010**), and more recently as a constellation of a few discrete functional groups (e.g., **Wieder et al. 2015**) or even of continuous hypothetical individuals using a trait-based approach (**Wang and Allison 2022**) in a structurally explicit soil environment (**Wang et al. 2019**). Echoing this trend of effectively embracing ecological complexity, we present empirical and theoretical bases for the TAM structure and demonstrate its impacts, arguably suggesting a move beyond the single fine-root pool paradigm in terrestrial ecosystem modelling, which, however, still needs to confront a myriad of uncertainties and challenges.

## 5. Uncertainties and challenges in realizing the TAM structure

Lingering uncertainties and challenges are closely tied to prevailing model assumptions as exemplified in the example realization. First, parameter uncertainties point us to a challenge of constraining partitioning while improving the representations of phenology and distribution in general. Second, going beyond the simplistic demand-driven nutrient acquisition assumption to integrate fine-root and mycorrhizal fungal functions with TAM will necessitate significant theoretical explorations and empirical investigations for model formulation, parameterization, and validation. However, to eventually realize TAM in ecologically realistic demography models (**Fig.2**), new challenges will emerge regarding age- and size-based changes in TAM that the big-leaf paradigm neglects, especially in ecosystems of high biological diversity.

### 5.1 A unified framework to simulate partitioning, phenology, and distribution

Partitioning, a key component in realizing a 3-pool TAM structure, is the most outstanding uncertainty. Sensitivity analyses (see **Methods** in the **Appendix**) show the parameters controlling partitioning are among the most sensitive ones (**Supplementary Fig. 3**). A direct reason for this sensitivity to partitioning is trait divergence from 1 to 3 pools with respect to, e.g., C/N, longevity, and chemical composition, as well as vertical differentiation in turnover time (**Table 1**). In addition to more constrained parameterizations of these traits, more accurate partitioning is essential for uncertainty reduction. Constraining the uncertainty of partitioning, however, is not as straightforward as simply improving parameterization of TAM traits, which can be informed by more field measurements. Instead, the interactive nature among partitioning, phenology, and vertical distribution, as indicated in **Section 3**, calls for a unified framework to simulate dynamic partitioning while accurately treating phenology and distribution to reduce the associated uncertainties.

Optimization techniques may offer such a unified framework, but it still faces a few immediate challenges. First, finding appropriate objectives for optimization with a balanced perspective of both plant and mycorrhizal fungi is non-trivial (e.g., **Bloom et al. 1985**; **Koide and Elliot 1989**). In addition, understanding of both phenology and distribution of TAM are in their infancy. Phenology is influenced by a plethora of factors including endogenous cues (**Joslin et al. 2001; Tierney et al. 2003**) and exogenous, abiotic cues (e.g., **Radville et al. 2016**), as well as microbial processes in the soil [see the review by **O’Brien et al. (2021)**]. This complex regulation is evident from mounting observations of variability in numbers of fine-root growth peaks (e.g., **Steinaker et al. 2010; McCormack et al. 2015b**) and of complex life history strategies of mycorrhizal fungi with spore dormancy (e.g., **Gianinazzi-Pearson et al., 1989; Bago et al. 2000**; **Defrenne et al. 2021**). TAM variability in phenology is far from being robust as it cannot yet account for all relevant abiotic factors. However, an independent treatment of TAM initiation and mortality allowing for asynchronous phenology between leaves and fine-root systems represents a substantial improvement. Still, the lack of asynchronous TAM phenology vertically means that the timing of partitioning at each depth is not fully accurate. Also, dynamic distribution, though constrained by dynamic nutrient availability in the soil profile, needs to account for other important factors (e.g., soil moisture and temperature). Addressing these interrelated processes under an optimization framework requires not only explicit seasonal observations of fine-root system dynamics across root orders and mycorrhizal fungi within soil profiles to disentangle environmental controls but requires explicit TAM functioning that goes beyond the demand-driven approach.

### 5.2 Connecting TAM with fine-root and mycorrhizal fungal functions

Relaxing the assumption of demand-driven nutrient uptake and competition entails an explicit formulation of the different functions of T, A and M pools and their interactions with soils. Incorporating explicit TAM functioning with respect to nutrient and water uptake will require explicit absorptive root and mycorrhizal fungi traits (**Table 1**) related to uptake of nutrients in various forms of nitrogen and phosphorous (e.g., **Yang et al. 2014; 2019**) and water (e.g., **Polverigiani et al. 2011; Jackisch et al. 2020; Kakouridis et al. 2020; Mackay et al. 2020).** Notably, mycorrhizal fungi in TAM, especially ectomycorrhiza and ericoid mycorrhiza, can directly mediate litter and soil organic matter decomposition by exuding enzymes and taking up organic nitrogen (e.g., **Gadgil and Gadgil 1971; Frey 2019**). Therefore, once TAM incorporates explicit root and mycorrhizal functions, it becomes highly necessary to make a concomitant change to microbially-explicit organic matter decomposition with exoenzymes instead of the first-order decomposition assumed in ELM. A microbially explicit decomposition model then needs to accommodate fine-root system exudates while handling explicit litter decomposition arising from the explicit turnover of TAM components, especially with fungal necromass (e.g., **Matamala et al, 2003; Strand et al. 2008; Fernandez et al. 2013; Sun et al. 2018**). These aspects can be addressed by combining a microbially explicit organic matter decomposition model interacting with the functioning of fine-root systems (e.g., **Sulman et al. 2018; Wang and Allison 2019**) with the existing approach of resolving plant-microbe competition for nutrient uptake based on the Equilibrium Chemistry Approximation theory (**Zhu et al. 2017; Wang and Allison 2019**). However, to capture the variability in these processes and their interactions arising from biological diversity, we need to move beyond the big-leaf paradigm.

### 5.3 Realizing TAM in demography models

Moving forward from the big-leaf structure to realizing TAM in demography models will require creative efforts to best capture patterns and processes belowground. The mosaic of forested landscapes so far has only been emphasized aboveground, and while previous studies have documented high heterogeneity in belowground processes, our example realization of TAM using the big-leaf approach circumvents this challenge (**Fig.2**). One outstanding issue towards this end is to simulate age- and size-related changes in TAM in the context of complete life cycles of growth, mortality, and reproduction for many different individuals under disturbances. For instance, evidence has indicated roles of mycorrhizal fungi change with plant community succession stage (e.g., **Pankow et al. 1991; Read 1991; Nara 2006**). One strategy of simulating age- and size-related changes would be conditioning TAM on a dynamic rooting depth of coarse roots (constrained by, for example, maximum rooting depth; **Fig.1; Table 1**), instead of using a constant value as widely assumed in current models. This change in rooting depth should involve geometric and allometric relationships with aboveground structural tissues (e.g., **Eshel and Grünzweig 2013; Brum et al. 2019; Tumber-Dávila et al. 2022**) while accounting for direct influences from, for example, soil hydrological conditions (e.g., **Stone and Kalisz 1991**; **Fan et al. 2007**). Addressing these aspects requires long-term observations of successional dynamics of not only the aboveground but also the belowground (e.g., **Rees et al. 2001;** “**We must get a grip on forest science — before it’s too late”, 2022**).

## 6. Implications of TAM for root ecology, ecosystem functioning, and ESMs

Confronting those above challenges holds promise to stimulate empirical ecology of fine-root systems, to drive theory-driven explorations of hypotheses of fine-root systems’ roles in ecosystem functioning, and to improve the prognostic capability of ESMs. First, adopting this structure would stimulate more targeted efforts at an increasing resolution of empirical research of fine roots and mycorrhizal fungi as discussed above on challenges (**Table 1**). The proposal of TAM is largely fueled by the belowground trait data revolution, especially with tremendous empirical progress in collecting function- and/or order-based fine-root traits including microbial associations (e.g., **McCormack et al. 2017; Iversen 2017; Iversen and McCormack 2017; Freschet et al. 2021**; **Tedersoo et al. 2021**). Building on standardized measurement protocols (**Freschet et al. 2021**), TAM may also stimulate further technical research and possible advances in, for example, neutron imaging (e.g., **Warren et al. 2013**), remote sensing (e.g., **Sousa et al. 2021**), and machine learning in image recognition (e.g., **Han et al. 2021**), to speed up characterization and classification of functional ‘roots’ within fine-root systems.

Undoubtedly, new measurements and causal relationships revealed by empiricists will improve formulation, parameterization, and validation of TAM in models of increasing complexity (**Fig. 2**). It is noteworthy that one of the major motivations for demography models is that local community-level processes are essential in modelling ecosystem functioning responding to various disturbances (e.g., **Wang et al. 2016; Fisher et al. 2018**). Increasing empirical and theoretical studies have indicated that root competition may even be more important than aboveground competition in shaping a plant community (e.g., **Gersani et al. 2001; Ljubotina and Cahill 2019; Cabal et al. 2020; Sauter et al. 2021**). With this explicit structure of fine-root systems in demography models of varying complexity, we can conduct theory-driven modelling studies to explore hypotheses of root-associated community-level processes underlying ecosystem functioning, which is otherwise challenging to experimentally track in the field.

We anticipate that all these empirical and theoretical advances arising from TAM eventually will contribute to improving ESMs both directly and indirectly. A more accurate and effective representation of fine-root systems across ecosystems may be directly incorporated into terrestrial ecosystem models with different vegetation structures, as implied by the example realization in temperate forest systems. Indirect benefits accompanying root improvement would also likely follow by helping to identify sources of uncertainty in components of aboveground and soil processes. One fundamental aspect might be to stimulate efforts to improve the scheme of global plant functional type classification by systematically integrating belowground traits (e.g., **Smith et al. 1997; Phillips et al. 2013**). This indirect avenue is particularly promising considering the historical dominance of top-down thinking in ESMs development (e.g., **Sellers et al. 1986**).

These direct and indirect effects together will improve the prognostic capability of ESMs to evaluate biosphere-atmosphere interactions in the Earth system.

## 7. Conclusions

Uncertain terrestrial ecosystem models still contain a huge data-model discrepancy with respect to fine-root system complexity. We speculate that closing this model-data gap will improve the prognostic capability of models across scales. To effectively simulate this complexity, we propose a function-based, 3-pool TAM structure representing transport and absorptive fine roots and mycorrhizal fungi to approximate the high-dimensional structural and functional variations within fine-root systems. Building upon theoretical and empirical bases of TAM as a balanced approximation between realism and simplicity, we quantitatively confirmed the significance of TAM for pools and fluxes of temperate forests arising from capturing the structural and functional differentiation within fine-root systems in a big-leaf land surface model. Though uncertainties and challenges remain towards realizing TAM in explicit demography models, TAM opens the door for more realistically and effectively capturing fine-root systems complexity underlying their functioning to eventually contribute to uncertainty reduction in ESMs.

## Acknowledgements

The Fine-Root Ecology Database (FRED), BW, CMI, DR, and XY were supported by the Biological and Environmental Research program within the Department of Energy’s Office of Science. The code of simple_ELM in Python is accessible at https://github.com/dmricciuto/simple_ELM/tree/rootcomplexity. Outputs from the simulations and code of analysis and visualization in Python are available at https://github.com/bioatmosphere/TAM. This research used resources of the Compute and Data Environment for Science (CADES) at the Oak Ridge National Laboratory, which is supported by the Office of Science of the U.S. Department of Energy under Contract No. DE-AC05- 00OR22725.

## Author Contributions

BW, MLM, DMR, XY, and CMI designed the research. BW performed the research, analyzed data, and wrote the first manuscript draft. MLM, DMR, XY, and CMI contributed to results interpretation and manuscript editing.

## Data Availability

The code of simple_ELM in Python is accessible at https://github.com/dmricciuto/simple_ELM/tree/rootcomplexity. Outputs from the simulations and code of analysis and visualization in Python are available at https://github.com/bioatmosphere/TAM. The database FRED is accessible at https://roots.ornl.gov/.

## 1. Methods

### 1.1 Simple ELM (sELM)

sELM is a simplified version of the Energy Exascale Earth System Model (E3SM) Land Model ELM, a state-of-the-art land surface model developed by Department of Energy, United States (**Golaz et al. 2019; Burrows et al. 2020**). ELM is traced back to the original CLM4.5 (**Oleson et al. 2013)** and the later version E3SM Land Model (ELM). sELM simplifies ELM by keeping only essential components simulating natural ecosystems in North America including vegetation and soil biogeochemistry (**Lu and Ricciuto 2019**). Following the same global PFT classification scheme as ELM, vegetation has the same structural pools (with a single fine-root pool) as ELM using the big-leaf approach. Although photosynthesis and GPP are replaced by a neural network surrogate built from ELM simulations, the whole scheme of allocation of photosynthates remains the same. Soil system, except for the lack of hydrology, is the same as the ELM with a 10-layer profile of 3.8018 m deep. Litter and soil organic matter decomposition follow the CTC (Converging Trophic Cascade) framework (**Oleson et al. 2013; Burrows et al. 2020**). Plant-microbe competition for mineral nitrogen is achieved via the relative demand approach without further differentiating between nitrogen forms (i.e., NH4 and NO3). Such a simplified model, in a more formal sense, is a high-fidelity mechanistic surrogate; by omitting irrelevant processes but keeping all essential components, it eases modifications of structures and processes and reduces computational burden for analyses of uncertainty and sensitivity, contributing to an acceleration of the development-evaluation-application cycle for Earth System Models in particular.

### 1.2 Realization of TAM

#### 1.2.1 Partitioning

We directly introduce three parameters (***frootpar_i***, where *i* indexes the three fine root pools of T, A, and M) to determine the spreading of allocation to a fine-root system (from either photosynthates or storage) among TAM pools. This approach simplifies the partitioning without considering explicitly the processes of movement of carbohydrates across the branching structure. With known partitioning (***frootpar_i***) and fine root C/N (***frootcn_i***), allocation allometry (ratio of allocation to total new growth to allocation to new leaf, *Callom/Nallom*) can be determined following:

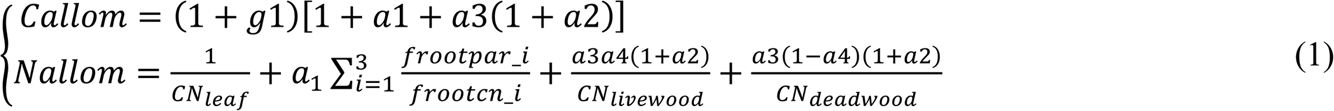

where ***a1*** through ***a4*** are allometric parameters that relate allocation between various tissue types, among which ***a1*** is the ratio of new fine root to new leaf carbon allocation (i.e., ***froot_leaf***), ***a2*** ratio of new coarse root to new stem carbon allocation, ***a3*** ratio of new stem to new leaf carbon allocation, ***a4*** ratio new live wood to new total wood allocation, and ***g1*** allocation ratio of growth respiration carbon to new growth carbon, as well as C/N of both live wood and dead wood. TAM pools are parameterized with different C/N ratios based on consistent observations of descending C/N while fine-root systems branching out (e.g., **Pregitzer et al. 1997; McCormack et al. 2015**).

#### 1.2.2 Phenology

In ELM, fine-root production is temporally controlled by leaf phenology in moving carbon from two direct sources: recent photosynthate and storage, making leaf and root production perfectly synchronous. These two sources, though ultimately originating from photosynthesis, can also be referred to as direct and indirect allocation, respectively. The allocation from leaf photosynthates to the fine-root system is based on an allometric relationship between leaves and fine roots (***froot_leaf*** = 1). This allocation is divided between direct allocation to the currently displayed fine-root system (***fcur***) and a storage pool to supply future growth for the whole fine-root system (i.e., indirect allocation; 1 - ***fcur***). This storage-based indirect allocation is also controlled by leaf phenology. Such a totally synchronous treatment between leaf and fine-root production is the case across all current ecosystem models, which, however, does not reflect the increasingly rich evidence that shoot and root phenology are not synchronous (e.g., **Perry 1971**; **Steinaker and Wilson 2008**; **Steinaker et al. 2010**; **McCormack et al. 2015b; Abramoff & Finzi 2015; Radville et al. 2016**). We make the following 3-step changes to decouple the phenological control of shoot and TAM in both deciduous and evergreen systems.

First, we make evergreen PFTs have the two-source structure of storage and recent photosynthate. In the current ELM, this two-source structure only applies to deciduous PFTs. Evergreen PFTs do not accumulate carbon and nitrogen in the storage pools for fine-root systems; instead, root production is solely determined by direct allocation of new photosynthates (i.e., ***fcur* = 1**), an assumption that lacks empirical support (e.g., **Huang et al. 2021**). Therefore, this treatment is changed for evergreen PFTs to have the same two-source structure as deciduous PFTs.

This structure enables storage-based production of (or indirect allocation to) fine-root systems controlled by TAM phenology.

Then, we decouple TAM phenology from leaf phenology for both deciduous and evergreen PFTs (**Fig.1**). In the current ELM, storage-based initiation of fine roots is synchronous with leaf-based control of unfolding (triggered by a growing degree threshold) and shedding (triggered by a critical day length). This synchronous treatment is changed to allow TAM initiation to behave independently in both deciduous and evergreen systems, which are determined by ***GDDsum*** (growing degree days with a 0-degree base) relative to a threshold value, ***GDDfroot_sum_crit***:

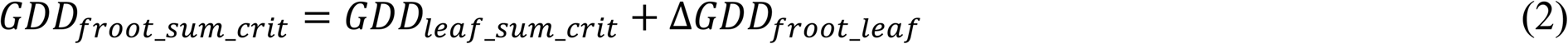

where ***ΔGDD_leaf_froot_*** is the difference between leaf and fine root threshold of GDD. Once ***GDDsum > GDD_froot_sum_crit_***, TAM initiation is triggered, and fluxes of carbon and nitrogen begin to move from storage pools (*CS_leaf_froot_*) to the fine-root system. This occurs via transfer pools (*CF_froot_stor,froot_xfer_*) calculated following:

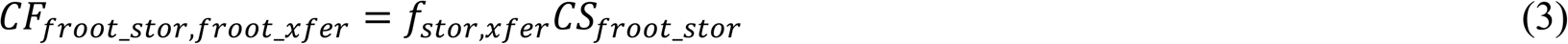

where *f_stor,xfer_* = 0.5, the fraction of current storage pool moved into the transfer pool for display over the incipient onset period (**Oleson et al. 2013**).

Finally, we make TAM turnover of deciduous PFTs dependent on pool-specific mortality.

For deciduous PFTs in the current ELM, the shedding of fine roots is synchronous with leaf shedding, which lacks empirical support. This synchronous treatment is thus changed to allow deciduous fine-root mortality to behave independently in deciduous systems in the same way as the fine roots of evergreen PFTs. See **Section 1.2.4** on TAM mortality.

#### 1.2.3 Distribution

In ELM and other land surface models, the prevailing option assumes that fine roots and coarse roots follow the same distribution as determined by prescribed fixed rooting and soil depths following:

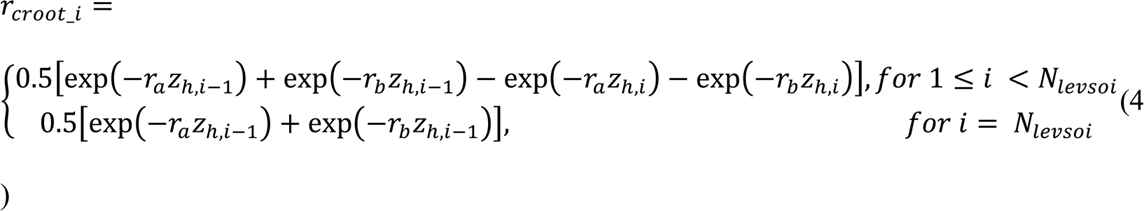

where ***ra*** and ***rb*** are PFT-dependent root distribution parameters. Because of the dynamic nature of fine-root systems, the vertical profile is modelled dynamically by assuming horizontal homogenization of TAM (that is, the same distribution profile across T, A, and M) constrained by nutrient and water availability:

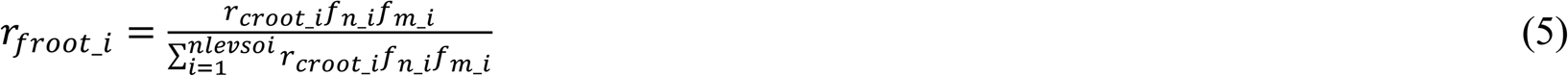

where ***fn_i*** and ***fm_i*** are the relative availability of nutrients and the relative availability of water at soil layer i, respectively. This nutrient constraint means that relatively more of the allocation goes to the depth where the relative nutrient availability is high; that is, a plant can direct allocation and distribution to the place where the nutrients are relatively richer. Note that root-water interactions are beyond the scope of this study and are not included in the analysis below. Such a realization of a dynamic profile contingent on a fixed coarse root profile partly accounts for the notion that coarser roots provide a structural constraint for potential fine-root system distribution dictated by the hierarchical branching structure of root systems.

#### 1.2.4 Mortality and litter input

With a 3-pool structure, turnover of the fine-root system is modelled explicitly in a vertically resolved way by introducing a mortality term. Mortality of fine roots and mycorrhizal fungi is endogenously controlled but subject to exogenous influences (e.g., **Eissenstat & Yanai 1997; Hendrick and Pregitzer 1997**). Therefore, consistent with the current approach in ELM modelling turnover using a mortality parameter in general, TAM mortality in each time step is determined by longevity distinguished among T, A, and M (***froot_long***) and constrained by a depth-correction term (**α**) as follows:

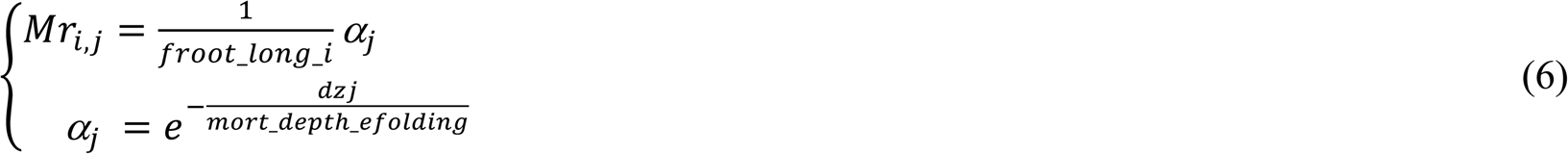

where ***froot_long_i*** is longevity of the three TAM pools near the soil surface, ***mort_depth_efolding*** is a constant of mortality rate decrease with depth, and ***dzj*** is depth of soil layer *j*.

Mortality of the T, A, and M pools are parameterized with different but decreasing lifespans based on relatively easy-to-make observations in shallow soil depths (**Supplementary Table 1**). This pattern is based on field observations of a dramatic increase in survivorship with diameter even within a very narrow range (e.g., ≤ 0.5 mm) (e.g., **Reid et al., 1993**; **Hendrick & Pregitzer 1993; Tierney & Fahey 2002; Edwards et al. 2004; Joslin et al. 2006; Espeleta et al. 2009**; **Riley et al. 2009; Xia et al. 2010**) and direct observations of lifespan and turnover in fungal tissues (**Allen and Kitajima 2013; McCormack et al. 2010**). The depth-correction term (***α***) further corrects the fine root and fungi longevity (***froot_long***) for increasing lifespan with soil depth (e.g., **Baddeley and Waston 2005; Iversen et al. 2008; McCormack et al. 2012; Germon et al 2015; Gu et al. 2017)**. This depth-based correction partly accounts for the plasticity of fine-root systems in response to vertical variability in soil resource availability (e.g., **Pregitzer et al. 1993; Eissenstat and Yanai, 1997**).

An explicit turnover of fine-root systems means explicit production of litters from the three pools. The TAM structure is coupled with existing litter decomposition model simply by aggregating litters across the 3 pools and depositing them in the three vertically resolved soil litter pools—labile, cellulose/hemicellulose, and lignin (**Oleson et al. 2013**). Each of the three fine-root pools is parameterized with fractions of these three different groups in terms of C (**Supplementary Table 1**). Nitrogen fluxes to the three litter pools from each of the three fine root pools are determined using the same fractions as used for carbon fluxes and their C/N ratios.

Additionally, maintenance and growth respiration follow the current ELM implementation (**Oleson et al. 2013**). Maintenance respiration rate is related to temperature and tissue N content without further differentiating intrinsic respiration rate and temperature sensitivity among TAM roots; varying N content differentiates respiration among TAM roots. Growth respiration is still calculated as a factor of 0.3 (***grperc***) times the total carbon in new growth including the fine-root system.

### 1.3 Parameterization and simulation

We implemented the changes as detailed above to realize the TAM structure in sELM. This implementation is referred to as TAM model (or TAM in short), while the original version is referred to as ELM. For an illustration purpose, we parameterized TAM and ELM for 2 temperate forests across the East US: Howland Forest (https://ameriflux.lbl.gov/sites/siteinfo/US-Ho1;

**Hollinger et al. 2021;** a single temperate evergreen needleleaf PFT) and Ozark (https://ameriflux.lbl.gov/sites/siteinfo/US-MOz; a single temperate deciduous broadleaf PFT; **Gu et al. 2016**). We used FRED (v3.0), a global Fine-Root Ecology Database to address below-ground challenges in plant ecology (https://roots.ornl.gov/; **Iversen et al. 2017; Iversen and McCormack 2021**), FunFun, a fungal trait database (**Zanne et al. 2020**), and published literature to derive PFT-specific parameter values/ranges for TAM (**Supplementary Table 1**). Simulation at each site was driven by forcings including daily radiation, maximum and minimum temperature, precipitation, day length, and a constant CO_2_ level (cycled 6 times).

### 1.4 Uncertainty and sensitivity analysis

TAM was compared with ELM (with default parameterization) by accounting for structural and parameter uncertainties introduced by the novel 3-pool structure. These comparisons involved outputs of interest including fluxes of GPP and heterotrophic respiration and pools of leaf and fine-root system TAM, as well as the nitrogen limitation factors for plant growth (i.e., Fraction of Potential Growth, *FPG = F_plant_uptake_/F_plant_demand_*_)_.

First, one case of preserving the fine-root system C/N of ELM in TAM was examined to see impacts of this structural shift and its associated uncertainties. To preserve the fine-root system

C/N in TAM (i.e., the default parameterization of C/N = 42), two specific scenarios were derived by solving the following equation, **Eq.7**, conditioned on the observed pattern of ascending C/N across the hierarchical branching orders within fine-root systems (e.g., **McCormack et al. 2015a**) and on prescribing one case of descending allocation from T through A (**TAM_descend**) and one case of ascending allocation (**TAM_ascend**):

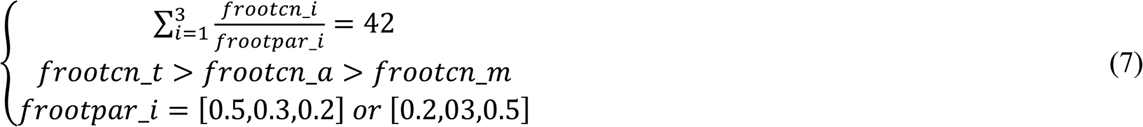

By prescribing two alternatives of partitioning fraction vector, the equation has two corresponding closed solutions of ***frootcn_i*:** [60,42,24] or [72,42,36]. In other words, by preserving the fine-root system C/N and keeping the same phenology and distribution between ELM and TAM, the uncertainty of the TAM structure for these two scenarios potentially arose from parameters of only chemistry and longevity. On top of these two cases, a structural uncertainty arising from nitrogen- constrained dynamic vertical distribution of new allocation was also examined, which were referred to as **TAM_descend_dd** and **TAM_ascend_dd**, respectively.

Next, without preserving the C/N ratio, a full implementation of TAM (including phenology) was compared with ELM. This comparison, relative to the above case, accounted for uncertainty arising from parameters including C/N and partitioning (6 parameters) and phenology-related parameters [3 parameters: ***gdd_crit*** (for evergreen PFT only)***, gdd_crit_gap, fcur***] **(Supplementary Table 1)**. That is, varying the partitioning and C/N parameters freely and phenology-related parameters under a full implementation of the proposed implementation of TAM, changes relative to ELM and their uncertainty were attributed to these newly introduced parameters.

Finally, a global sensitivity analysis of TAM was performed with 8 (7 for evergreen PFT which does not have ***crit_dayl***) more existing parameters in ELM (**Supplementary Table 2**). All parameter uncertainty was quantified by 1080 ensemble runs from Monte-Carlo sampling from parameters’ uniform uncertainty ranges and uncertainty propagation through the model. Attribution of TAM uncertainty was then achieved via variance-based sensitivity analysis with Sobol’s sensitivity indices (**Sobol 2003; Ricciuto et al. 2018**). The Sobol’s sensitivity indices were indirectly derived from a polynomial chaos expansion of the outputs via a new Weighted Iterative Bayesian Compressive Sensing (WIBCS) algorithm using the Uncertainty Quantification Toolkit (**UQTk v3.0.2**; https://github.com/sandialabs/UQTk).

## 2. Results

**Supplementary Table 1.**
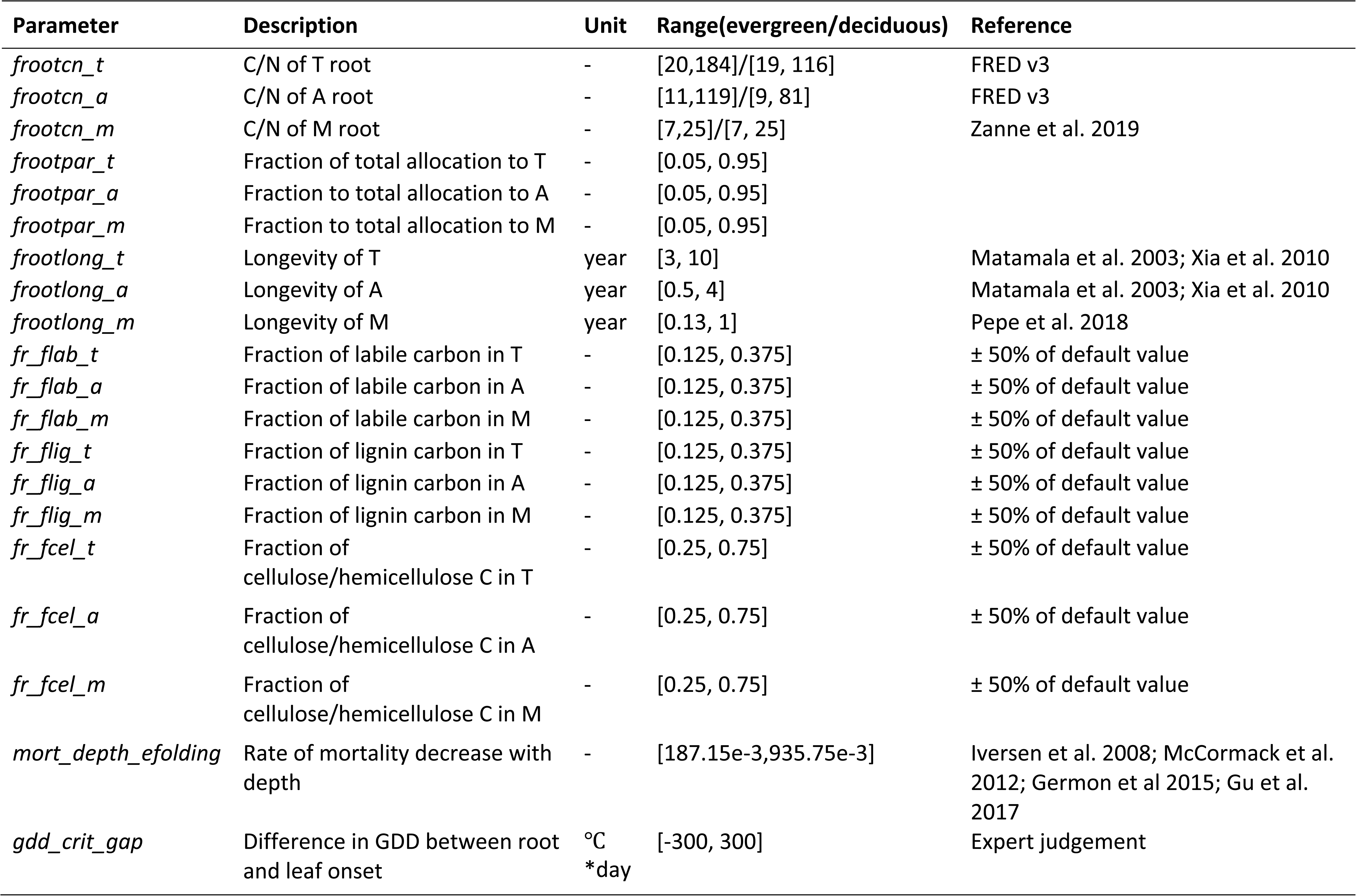
List of introduced parameters and their uncertainty ranges.

**Supplementary Table 2.** List of existing parameters of interest in ELM and their uncertainty ranges.

| Parameter | Description | Unit | Range(evergreen/deciduous) |
| --- | --- | --- | --- |
| *gdd_crit* | Leaf onset GDD | °C *day | 150, 750 |
| *fcur* | fraction of root allocation to display in current time step | - | [0.5, 1.0]/[0, 1] |
| *froot_leaf* | ratio of new fine root to new leaf carbon allocation | - | [0.5, 1.5] |
| *slatop* | Specific leaf area at canopy top | m<sup>2</sup> gC<sup>-1</sup> | [0.005,0.015]/[0.015,0.045] |
| *crit_dayl* | Critical day length for leaf senescence in deciduous | Seconds | [36000, 43000] |
| *leafcn* | leaf C/N | - | [17.5, 52.5]/[12.5, 37.5] |
| *roota_par* | Rooting depth distribution parameter | m-1 | [1.5,4.5]/[3, 9] |
| *rootb_par* | Rooting depth distribution parameter | m-1 | [0.625, 1.875]/[1, 3] |
| *br_mr* | Base rate for maintenance respiration (MR) | umol m<sup>-2</sup> s<sup>-1</sup> | [1.26e-06, 3.78e-06] |
| *q10_mr* | Temperature sensitivity for MR | - | [0.75, 2.25] |

**Supplementary Fig. 1.**
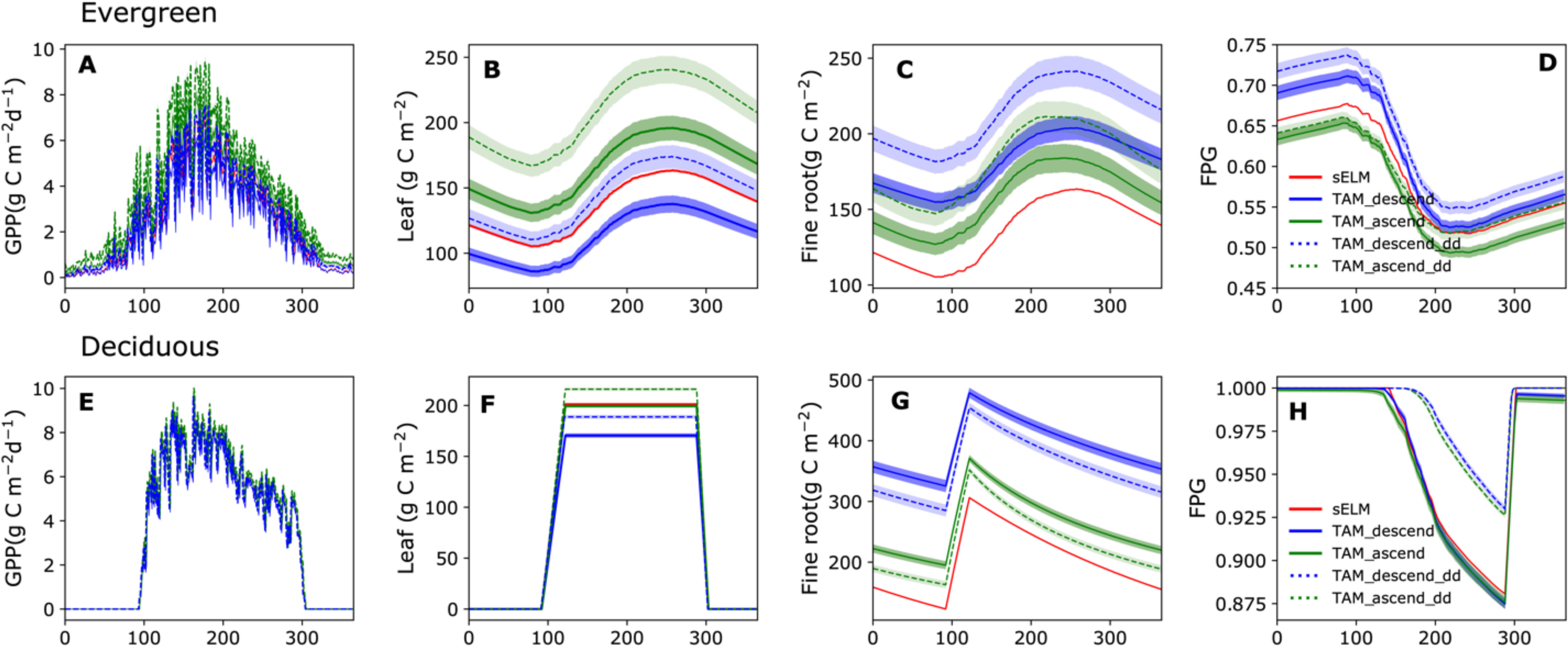
Seasonal dynamics of GPP, leaf mass, and fine-root mass, as well as FPG (Fraction of Potential Growth) of TAM against the 1-pool structure of sELM under preservation of bulk fine-root system stoichiometry.

**Supplementary Fig. 2.**
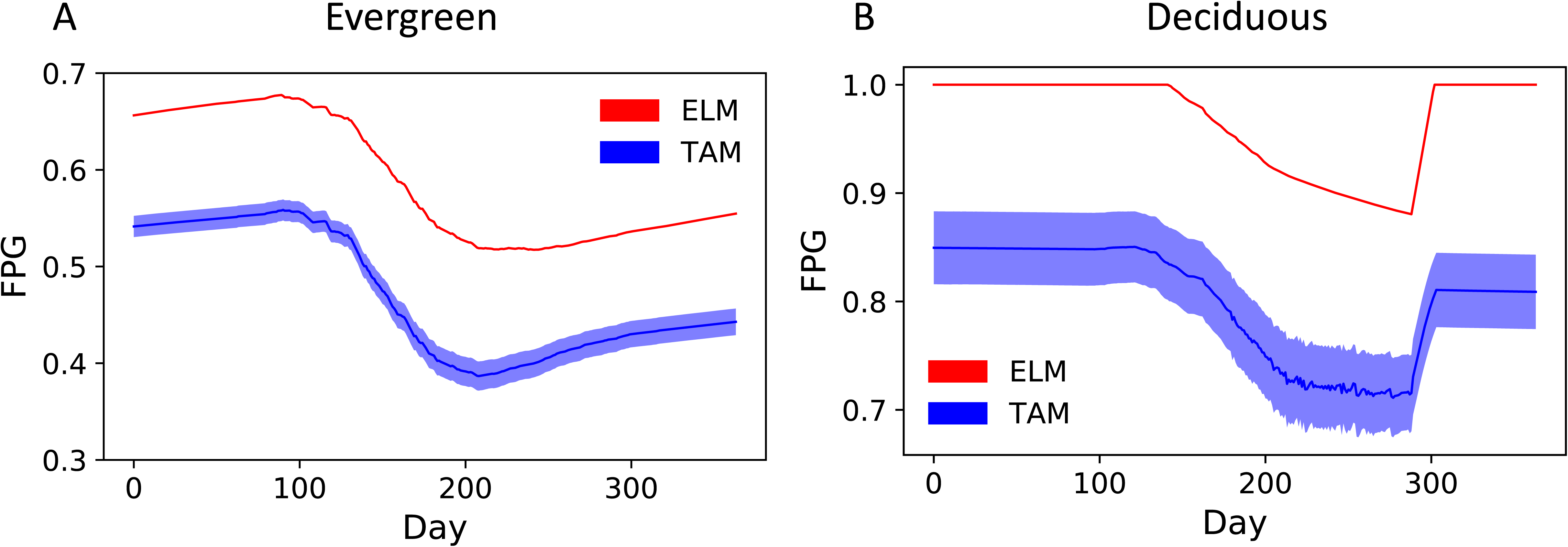
Seasonal variation in FPG (Fraction of Potential Growth) under a full implementation of TAM corresponding to **Fig.4** in the main text.

**Supplementary Fig. 3.**
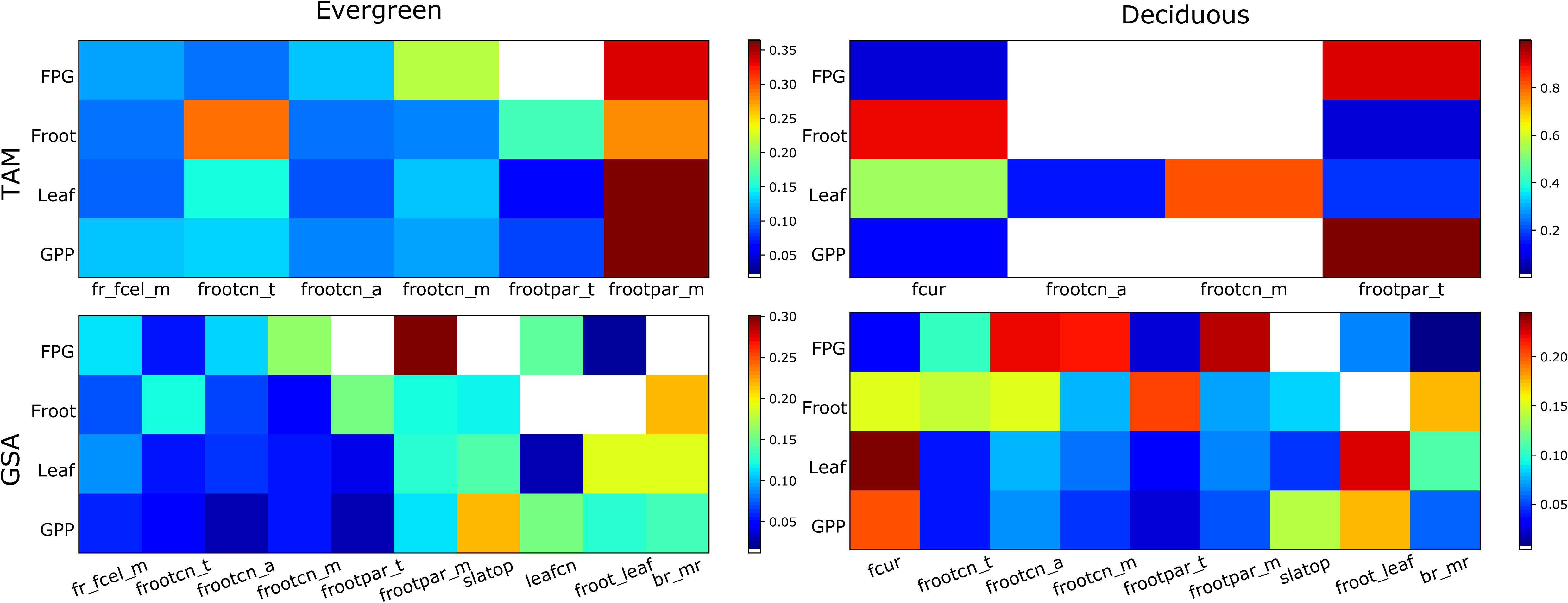
Attribution of TAM and global uncertainty with the main effect sensitivity index for the two sites. The TAM sensitivity corresponds to **Fig.3** in the main text. The global sensitivity analysis also includes some of existing parameters (**Supplementary** Table 2). Only the most relevant input parameters for each forest site are shown. The color code for each row is scaled according to the highest contributor to improve visibility.

## Notes

https://github.com/bioatmosphere/TAM

### Competing Interest Statement

The authors have declared no competing interest.

